# Sporulation-specific cell division defects in *ylmE* mutants of *Streptomyces coelicolor* are rescued by additional deletion of *ylmD*

**DOI:** 10.1101/214072

**Authors:** Le Zhang, Joost Willemse, Paul A. Hoskisson, Gilles P. van Wezel

## Abstract

Cell division during the reproductive phase of the *Streptomyces* life-cycle requires tight coordination between synchronous formation of multiple septa and DNA segregation. One remarkable difference with most other bacterial systems is that cell division in *Streptomyces* is positively controlled by the recruitment of FtsZ by SsgB. Here we show that deletion of *ylmD* (SCO2081) or *ylmE* (SCO2080), which lie in operon with *ftsZ* in the *dcw* cluster of actinomycetes, has major consequences for sporulation-specific cell division in *Streptomyces coelicolor*. Electron and fluorescence microscopy demonstrated that *ylmE* mutants have a highly aberrant phenotype with defective septum synthesis, and produce very few spores with low viability and high heat sensitivity. FtsZ-ring formation was also highly disturbed in *ylmE* mutants. Deletion of *ylmD* had a far less severe effect on sporulation. Interestingly, the additional deletion of *ylmD* restored sporulation to the *ylmE* null mutant. YlmD and YlmE are not part of the divisome, but instead localize diffusely in aerial hyphae, with differential intensity throughout the sporogenic part of the hyphae. Taken together, our work reveals a function for YlmD and YlmE in the control of sporulation-specific cell division in *S. coelicolor*, whereby the presence of YlmD alone results in major developmental defects.

## INTRODUCTION

In unicellular bacteria, cell division divides a mother cell in two identical daughter cells, each containing a single copy of the chromosome. The control of cell division thereby revolves around finding the mid-cell position, and chromosome segregation and septum synthesis are closely coordinated in time and space to avoid DNA damage by the nascent septum. The cell division scaffold is formed by FtsZ, which is a homologue of tubulin ^1^ and forms a contractile ring (or Z-ring) that mediates the recruitment of the cell division machinery to the division site (reviewed in ^2,3^). Septum-site selection and Z-ring stabilization are mediated by proteins like FtsA and ZipA ^4–6^, ZapA ^7^ and SepF ^8,9^. Z-ring (dis-)assembly is thereby actively controlled (reviewed in ^10^).

Streptomycetes are filamentous Gram-positive bacteria that belong to the phylum of Actinobacteria. These bacteria produce over 60% of all known antibiotics and many other bioactive natural products ^11,12^. Exponential growth of the vegetative hyphae is achieved by apical growth and branching. At this stage of the life cycle, cell division does not affect physical separation of the cells, but instead long syncytial cells are formed that are separated by cross-walls ^13^. Hence, streptomycetes are model organisms for the study of multicellularity and bacterial morphogenesis ^14,15^.

Most divisome components except FtsZ are not required for vegetative division, presumably reflecting the lack of constriction ^16^. Spacing between the cross-walls is highly variable, and little is known of the way septum-site selection is controlled. Recently, using cryoelectron tomography, our lab and others showed that intracellular membrane assemblies or cross-membranes are involved in DNA protection during septum synthesis in young vegetative hyphae, suggesting a novel way of cell-division control ^17,18^. In addition, these multicellular bacteria have a complex cytoskeleton, which among others plays a role in the organization of the tip growth machinery ^19,20^.

Canonical division resulting in cell fission occurs during sporulation-specific cell division, which requires all components of the divisome ^16,21^. At this stage of the life cycle up to a hundred septa are formed in a short time span, following a highly complex process of coordinated cell division and DNA segregation, and visualized as long ladders of Z-rings ^22^. Eventually, chains of unigenomic spores are formed, which have a thick protective spore wall facilitating long-term survival in the environment. Though *ftsZ* null mutants are viable, they fail to produce septa and hence do not sporulate, cell division is not essential for growth of *Streptomyces*, which provides a unique system for the study of this process ^23,24^.

Sporulation-specific cell division is controlled by the SsgA-like proteins (SALPs), which are exclusively found in morphologically complex actinobacteria ^25,26^. The canonical view is that cell division is negatively controlled by the action of the Min system that inhibits division away from midcell, and by nucleoid occlusion (Noc) to avoid septum synthesis near the chromosome to avoid DNA damage, as seen in *B. subtilis* and *E. coli*. In contrast, cell division in streptomycetes is positively controlled by the recruitment of FtsZ to future septum sites by SsgB, in an SsgA-dependent manner ^27^. As a consequence, both SsgA and SsgB are required for sporulation ^28,29^. We recently showed that SepG (formerly called YlmG) assists in docking of SsgB to the membrane, and also plays a major role in maintaining the nucleoid shape in the spores ^30^.

Many of the genes for the components of the divisome and the cell-wall biosynthetic machinery are located in the so-called *dcw* cluster (division and cell-wall synthesis; ^31^). Most of these genes have been studied extensively and their functions have been well characterized. However, little is known of the genes *ylmD* and *ylmE* that lie immediately downstream of, and likely form an operon with, *ftsZ* on the genome of streptomycetes and many other bacteria, including firmicutes. Earlier work on a mutant of *S. venezuelae* lacking both *ylmD* and *ylmE* showed that the double mutant had no obvious phenotypic defects ^32^.

In this study, we show that deletion of *ylmE* alone results in severe cell division defects, while deletion of *ylmD* only slightly affects sporulation. Interestingly, the cell division defects of *ylmE* mutants were rescued by the additional deletion of *ylmD*, and *ylmDE* mutants had no obvious sporulation defects. These data strongly suggest that expression of YlmD alone is detrimental for sporulation-specific cell division in streptomycetes, which is counteracted by YlmE. This is consistent with the phylogenetic evidence that some bacteria only harbor an ortholog of *ylmE*, whereas *ylmD* never occurs without *ylmE*.

## RESULTS & DISCUSSION

### Phylogenetic analysis of YlmE (SCO2080) and YlmD (SCO2081)

Many of the genes in the *dcw* gene cluster have been extensively studied and their functions are well established. However, this is not the case for *ylmD* and *ylmE*, which lie immediately downstream of *ftsZ* in many bacteria. In all *Streptomyces* genomes analyzed, *ftsZ* (SCO2082), *ylmD* (SCO2081) and *ylmE* (SCO2080) form an operon, an observation that is supported by high-resolution transcript mapping ^33^; moreover there is apparent translational fusion between *ftsZ* and *ylmD* (overlapping start and stop codons) and only 6 nt spacing between *ylmD* and *ylmE*. This transcriptional coupling to *ftsZ* suggests that these genes may play a prominent role in the cell division process. Transcript levels of *ftsZ* are similar to those of *ylmD* and *ylmE* during vegetative growth on MM agar plates; however, transcription of *ftsZ* is enhanced during sporulation, while that of *ylmD* and *ylmE* is not significantly altered ^34^. YlmD and YlmE are wide-spread in Gram-positive bacteria, and particularly in actinobacteria. SCO2081 and SCO2080 share 32% aa identity with YlmD and YlmE of *B. subtilis*, respectively. Orthologues of these proteins are also found in several genera of Gram-negative bacteria. Phylogenetic analysis of the YlmD and YlmE proteins in actinobacteria shows that while YlmE is widespread, YlmD is often absent in actinobacteria, such as *Stackebrandtia, Catenulispora, Salinispora, Micromonospora, Amycolatopsis* and *Mycobacterium*, several of which are spore-forming actinobacteria (Fig. 1). Remarkably, there are no examples of *ylmD* being present in the absence of *ylmE* and thus it would appear that loss of YlmD (SCO2081) has occurred on multiple occasions, given that there is wide but patchy distribution of this gene across distinct actinobacteria lineages and the sequences clade tightly within the accepted actinomycete phylogenies.

**Figure 1.**
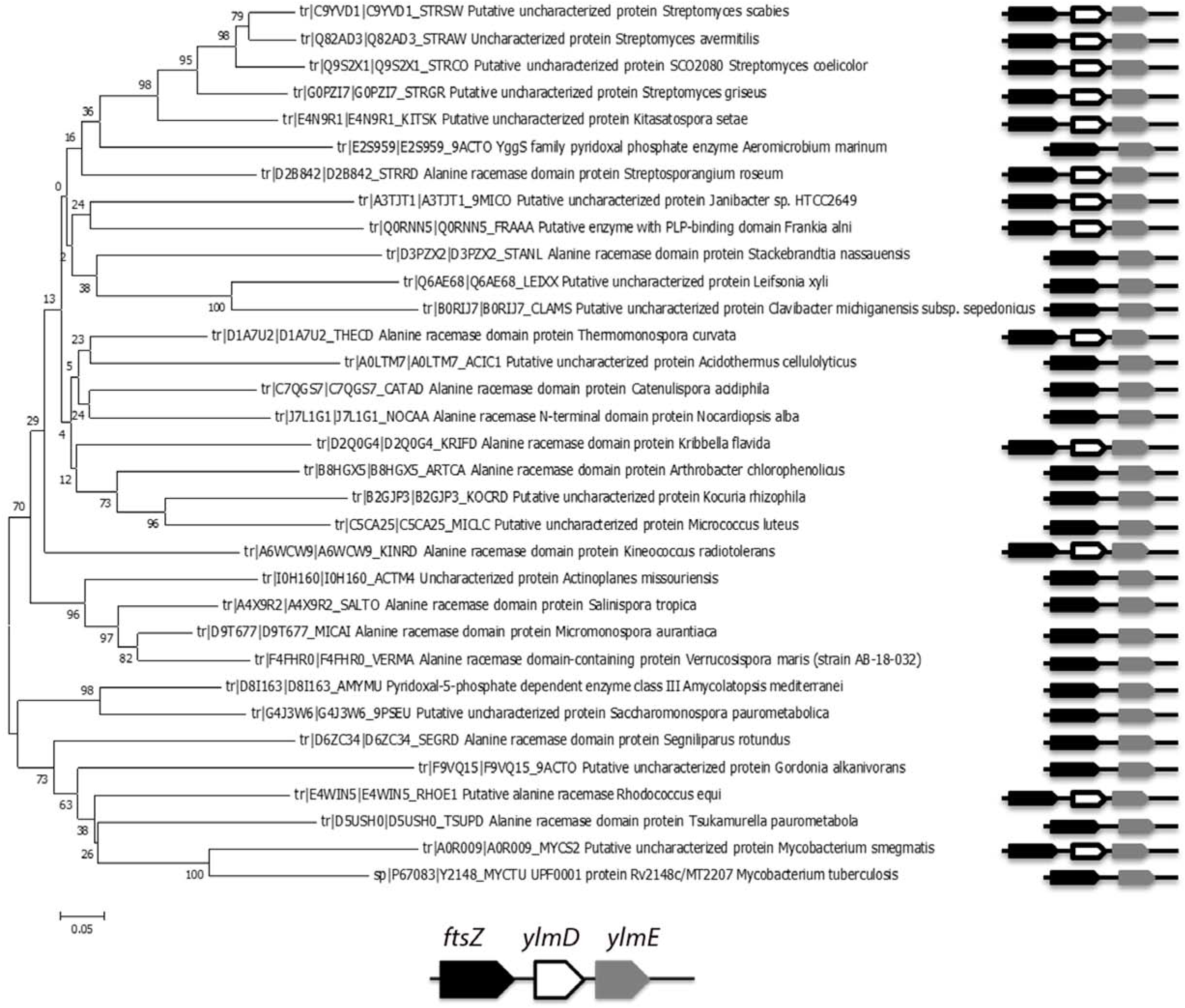
Phylogenetic analysis of YlmE in actinobacteria and its genetic linkage to *ylmD*. **A** phylogenetic tree is shown of YlmE in actinobacteria (left) and genetic linkage of *ylmE* (grey) to *ftsZ* (black) and *ylmD* (white) (right).

Analysis of *ylmD* using the EMBL String engine ^35^ shows functional linkage in two groups to cell division associated genes (*sepF* (SCO2079), SCO2085 and *ftsZ*) along with a group of mainly hypothetical proteins (STRING Data link: http://bit.ly/2gc5kCB). YlmD has a domain that has homology to the multiple-copper polyphenol oxidoreductase laccases, which are oxidoreductases that are widely distributed in both prokaryotes and eukaryotes ^36^. A similar analysis of YlmE also indicates a functional linkage to cell-wall biosynthesis and cell division (STRING Data link: http://bit.ly/2gbPyHK). YlmE is a member of the family of YBL036c-like proteins, which generally contain pyridoxal 5-phosphate dependent enzymes. The structure of YBL036c from *Saccharomyces cerevisiae* was resolved at 2.0 Å resolution (PDB 1CT5;^37^). The protein has homology to the N-terminal domains of alanine racemases but lacks of the β-sandwich domain which would likely limit the activity of YBL036c as alanine or non-specific amino acid racemase ^38^. To test the hypothesis that YlmE may have alanine racemase activity and thus may play a role in determining the amino acid composition of the peptidoglycan, we tested purified YlmE for alanine racemase assays as described previously ^39^, using alanine racemase Alr as positive control. Whereas purified Alr successfully catalyzed the conversion of L-alanine to D-alanine, YlmE could not perform this reaction under the same conditions, and over-expression of YlmE failed to restore a D-Ala prototrophy to *alr* null mutants (data not shown). These observations coupled with protein structure homology data make it highly unlikely that YlmE functions as an alanine racemase *in vivo*.

### *ylmD* and *ylmE* are required for proper sporulation

To analyze the role of *ylmD* and *ylmE*, deletion mutants were created in *S. coelicolor* M145 as detailed in the Materials and Methods section. The +25 to +696 region of *ylmE* (SCO2080) or the +25 to +705 region of *ylmD* (SCO2081) were replaced by the apramycin resistance cassette, followed by deletion of the cassette using the Cre recombinase so as to avoid polar effects. For each mutant, four independent mutants were selected and all had the same phenotype. Therefore, one was selected for more detailed analysis, designated GAL47 (*S. coelicolor* M145 Δ*ylmD*) and GAL48 (*S. coelicolor* M145 Δ*ylmE*). The M145 *ylmDE* mutant (GAL130) was created by replacing *ylmD* in *ylmE* null mutant GAL48 by the apramycin resistance cassette.

The *ylmD* null mutant GAL47 had a wild-type-like appearance, while *ylmE* null mutant GAL48 hardly produced any grey pigment after 5 days incubation on SFM agar plates, indicative of a failure to complete full development. Surprisingly, *ylmDE* mutant GAL130 had a grey appearance similar to that of the parental strain M145 (Fig. 2A). Phase-contrast light microscopy of impression prints of the top of the colonies demonstrated that while the parent produced typical long spore chains, the *ylmD* null mutant produced abundant but aberrantly sized spores, the *ylmE* null mutant produced fewer spores and those produced were unusually large. The *ylmDE* double mutant produced abundant spores, though some were irregularly sized (Fig. 2B and Fig. S1). The same defect in sporulation was observed on R5 and MM mannitol agar plates. Antibiotic production was not affected, and the mutants produced normal levels of the pigmented antibiotics actinorhodin and undecylprodigiosin.

**Figure 2.**
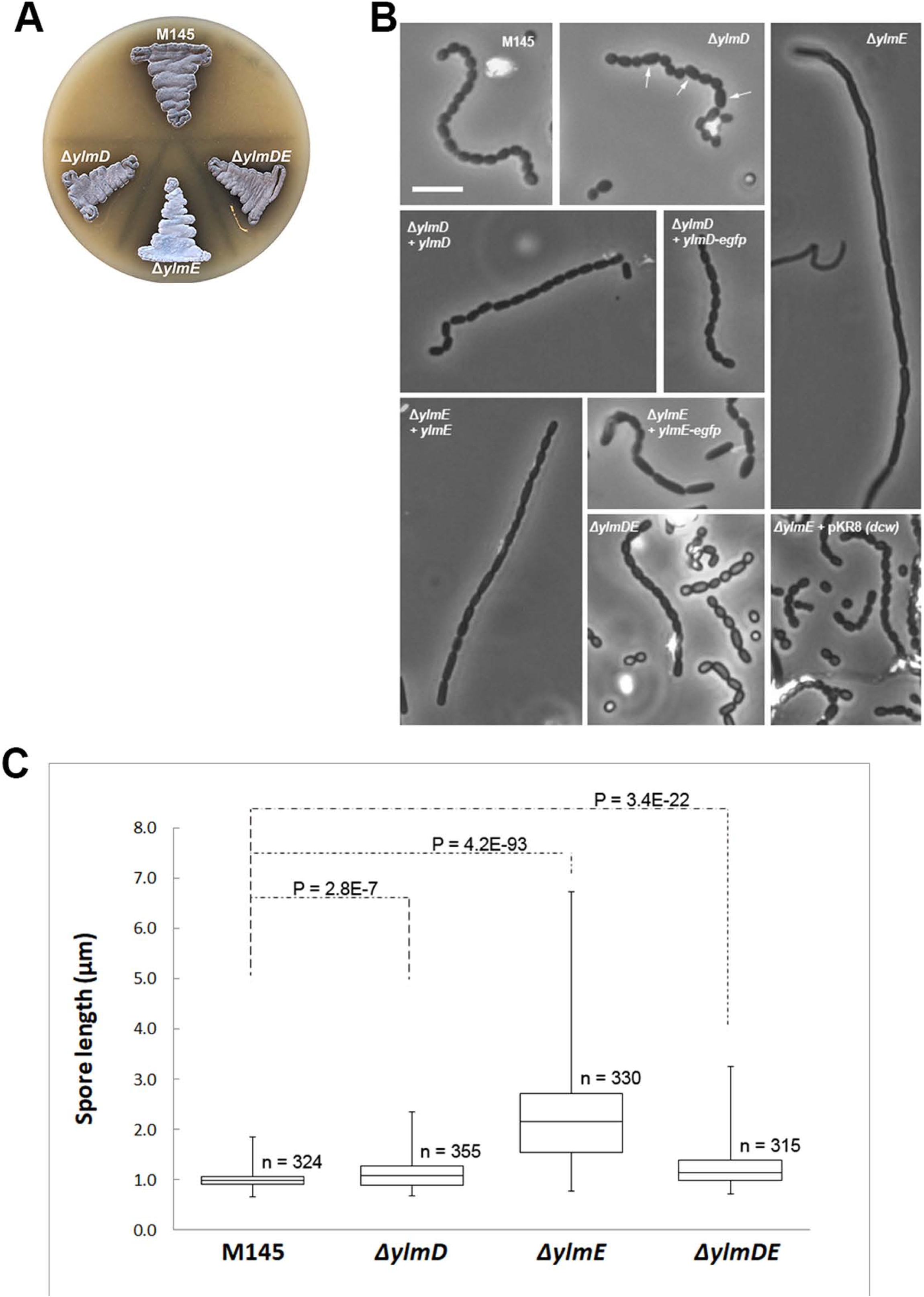
Phenotype of the *S. coelicolor ylmD* and *ylmE* mutants. (A) sporulation of *S. coelicolor* M145 and its *ylmD, ylmE* and *ylmDE* mutants on SFM agar after 5 days incubation. Note the lack of grey pigmentation of the *ylmE* mutant, indicative of a sporulation defect. (B) phase-contrast micrographs of impression prints of the strains shown in (A) as well as the complemented mutants. Arrows point at irregularly shaped spores in the *ylmD* mutant. The sporulation defect of the *ylmD* mutant could be complemented by introduction of wild-type *ylmD* and *ylmD-eGFP*, while complementation of the *ylmE* mutant by wild-type *ylmE* or *ylmE-egfp* restored sporulation, although irregularly sized spores were often produced. Full complementation of *ylmE* mutants was obtained by introduction of pKR8, which contains part of the *S. coelicolor dcw* cluster. Bar, 5 μm. (C) Size distributions of wild-type spores and those of *ylmD, ylmE* and *ylmDE* mutants presented as boxplots. Data are presented as median and interquartile range in boxplots, with whiskers spread to the maximal and minimal values. The numbers (n) of spores measured for each strain are indicated. Assessment of normality of spore length data was performed by a Kolgomorov-Smirnov test, the results showing that spore length data for all mutants followed non-normal distribution (*D*_max_ >*D*_crit_). The shown two-tailed P values between each mutant and the parental strain were calculated using a Mann-Whitney U test. The two-tailed P values were far below 0.05, which shows that the sizes of the mutant spores deviated significantly from those of wild-type spores.

To further ascertain that the sporulation defects were indeed solely due to the deletion of the respective genes, we introduced plasmids expressing *ylmD* or *ylmD-egfp* in the in the *ylmD* null mutant and *ylmE* or *ylmE-egfp* in the *ylmE* mutant, in all cases with transcription directed from the native *ftsZ* promoter region. While the majority of the spores of the complemented *ylmD* mutant had a regular appearance, those of the complemented *ylmE* mutants still showed variable lengths (Fig. 2B). This partial complementation suggests that the deletion of *ylmE* may have polar effect on the expression of its downstream genes. This is supported by the fact that introduction of a plasmid harboring the entire region from *murD* to *divIVA* fully restored sporulation to *ylmE* mutants (Fig 2B & Fig. S2). Promoter probing using the *redD* reporter system ^40^ showed that besides the intergenic promoter between *ylmE* and *sepF*, an additional promoter is located within *ylmE*, suggesting that deletion of the entire *ylmE* gene negatively affects the transcription of *sepF* (data not shown). This may explain why *ylmE* alone failed to fully complement the *ylmE* null mutant.

To quantify the spore lengths and their size distribution (*i.e*. variability), approximately 300 spores from the parental strain *S. coelicolor* M145, its *ylmD* and *ylmE* mutants and the genetically complemented strains were measured from light images from phase-contrast microscopy (100x magnification). Spore sizes were compared using a boxplot analysis (Fig. 2C). Wild-type spores typically had lengths between 0.8-1.2 μm, with an average length of 1.01 ± 0.15 μm, while *ylmD* mutant spores showed a slightly broader distribution (Fig. 2C), with an average length of 1.12 ± 0.27 μm. The average length was reduced to wild-type levels by complementation, namely to 1.03 μm and 1.05 μm by introduction of *ylmD* and *ylmD-egfp*, respectively. Spores of the *ylmE* null mutant were much larger and with a significantly wider variation, showing an average length of 2.23 ± 0.86 μm (Fig. 2C). Genetic complementation of the *ylmE* mutant restored spore length variation of *ylmE* mutant to wild-type levels, with an average size of 1.03 ± 0.19 μm. Partial complementation of the *ylmE* null mutant was seen when constructs expressing *ylmE* or *ylmE-egfp* were introduced into the mutant, with spore lengths of 1.62 ± 0.50 μm and 1.85 ± 0.59 μm, respectively. The average spore length of the combined *ylmDE* mutant was much closer to that of the parental strain, namely 1.23 ± 0.34 μm, thereby showing significantly smaller size variation than *ylmE* null mutants (Fig. 2C). The statistical validity of the variations in spore sizes between mutants and the parental strain was validated by a Kolgomorov-Smirnov test and a Mann-Whitney U test, which confirmed that the sizes of mutant spores deviated significantly from those of wild-type spores.

To study the spores of *ylmD* and *ylmE* null mutants at high resolution, cryo-scanning electron microscopy (SEM) was performed, which again demonstrated the sporulation defects. The parental strain *S. coelicolor* M145 produced abundant spore chains, with nearly all hyphae fully developed into mature spore chains (Fig. 3AB). The *ylmD* null mutant GAL47 frequently produced aberrantly sized spores (Fig. 3CD), while in *ylmE* null mutant GAL48, precious few and often aberrant spores were identified (Fig. 3EF). The same approach was used to create *ylmD* and *ylmE* mutants in *Streptomyces lividans* 66, and these again showed the same morphology, strongly supporting the notion that the observed phenotypes were solely due to the mutations in *ylmD* or *ylmE* (data not shown). The highly similar phenotypes of the *S. coelicolor* and *S. lividans* mutants support the notion that the genes play a key role in sporulation-specific cell division and are consistent with the conserved function of the genes in sporulation-specific cell division in *Streptomyces*.

**Figure 3.**
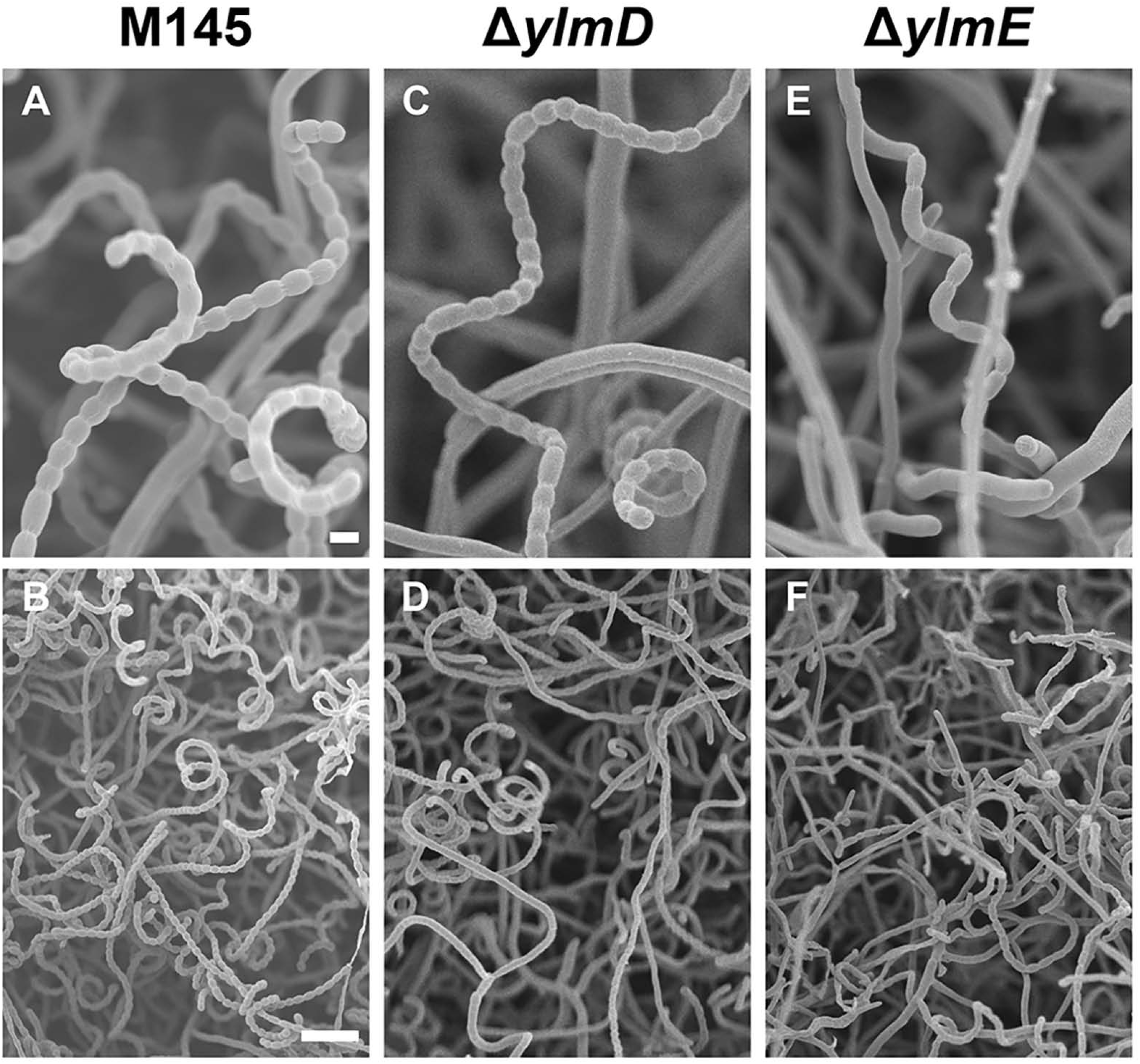
Cryo-scanning electron micrographs of aerial hyphae of *S. coelicolor* M145 and its *ylmD* and *ylmE* mutants. The parental strain produced wild-type spores, the *ylmD* mutant produced abundant but often irregular spores, while the *ylmE* null mutant produced occasional spores with highly irregular sizes. Cultures were grown on SFM agar plates for 5 days at 30°C. Bars: top row, 1 μm; bottom row, 5 μm.

### Deletion of *ylmD* and especially *ylmE* results in reduced robustness of spores

Viability of the spores was tested by plating around 1000 spores - counted in a hemocytometer - of *S. coelicolor* M145 and its mutants GAL47 (Δ*ylmD*) and GAL48 (Δ*ylmE*) onto SFM agar plates. While the wild-type strain had close to 100% viability, spores of the *ylmD* and *ylmE* mutants had reduced viability, with 50% and 60% viable spores, respectively. Spores of in particular the *ylmE* mutant were heat sensitive, with only 11 ± 1% viability after 15 min incubation at 58°C, as compared to 47 ± 6% survival for *ylmD* null mutant spores and 55 ± 5% for the parental strain. Genetic complementation of the *ylmE* null mutant restored wild-type survival to heat shock (58 ± 3%). The same was true when *ylmD* was deleted in the *ylmE* null mutant (56 ± 4% survival).

To analyze the possible cell-wall defects in more detail, all strains were grown on SFM agar plates for 5 days and analyzed by Transmission Electron Microscopy (TEM) (Fig. 4). The parent produced typical spore chains and thick-walled spores. Conversely, *ylmD* and *ylmE* mutant spores were deformed and highly irregular. In particular the spores of the *ylmE* mutant had a thin wall similar to that of hyphae, suggesting that spore-wall synthesis was compromised.

**Figure 4.**
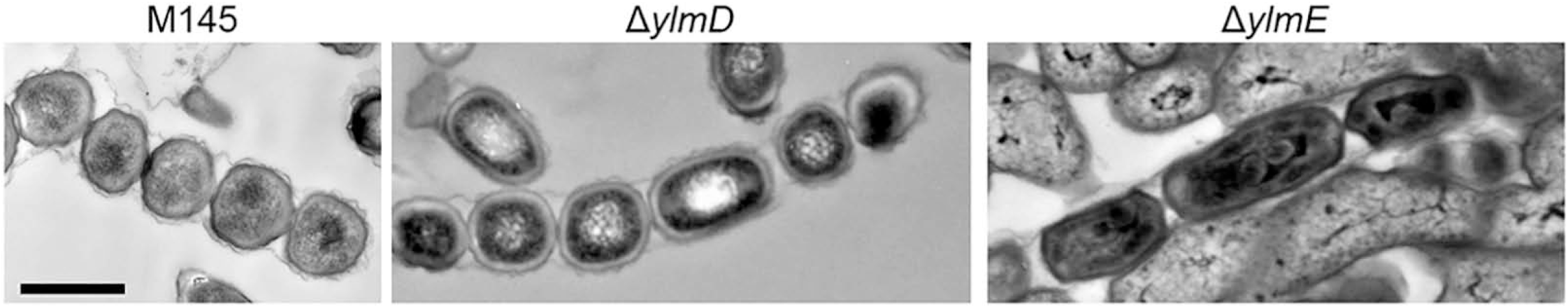
Transmission electron micrographs of spores of *S. coelicolor* M145 and its *ylmD* and *ylmE* mutants. Wild-type spores (M145) show regular sizes and appearance (A). In contrast, spores of the *ylmD* (B) and *ylmE* (C) null mutants have an irregular appearance. Note the lighter appearance of the spore wall in the *ylmD* null mutant and the lack of the typical thick spore wall in *ylmE* mutants. Cultures were grown on SFM agar plates for 5 days at 30°C. Bar, 500 nm.

### YlmD and YlmE are required for peptidoglycan synthesis at the septum

To analyze cell wall and membrane distribution in *ylmD* and *ylmE* mutants, fluorescence microscopy was performed on five-days old SFM surface-grown cultures of mutants GAL47 (M145 Δ*ylmD*) and GAL48 (M145 Δ*ylmE*) and compared to the parental strain M145. Peptidoglycan precursors were stained with FITC-WGA or Oregon-WGA and membranes visualized by staining with FM5-95. In wild-type pre-division aerial hyphae, long symmetrical septal ladders were observed when stained for cell wall or membranes (Fig. 5A). In contrast, aerial hyphae of the mutants showed highly disturbed cell wall and membrane distribution, which was more pronounced in *ylmE* than in *ylmD* mutants (Fig. 5A). Septation of *ylmD* mutants was complemented via re-introduction of *ylmD;* similarly, septation was restored to the *ylmE* mutant by re-introduction of *ylmE*, although peptidoglycan synthesis and septum spacing was still irregular (Fig. 5B). These data again show that the consequences of deleting *ylmE* are much more severe for sporulation-specific cell division then when *ylmD* is deleted.

**Figure 5.**
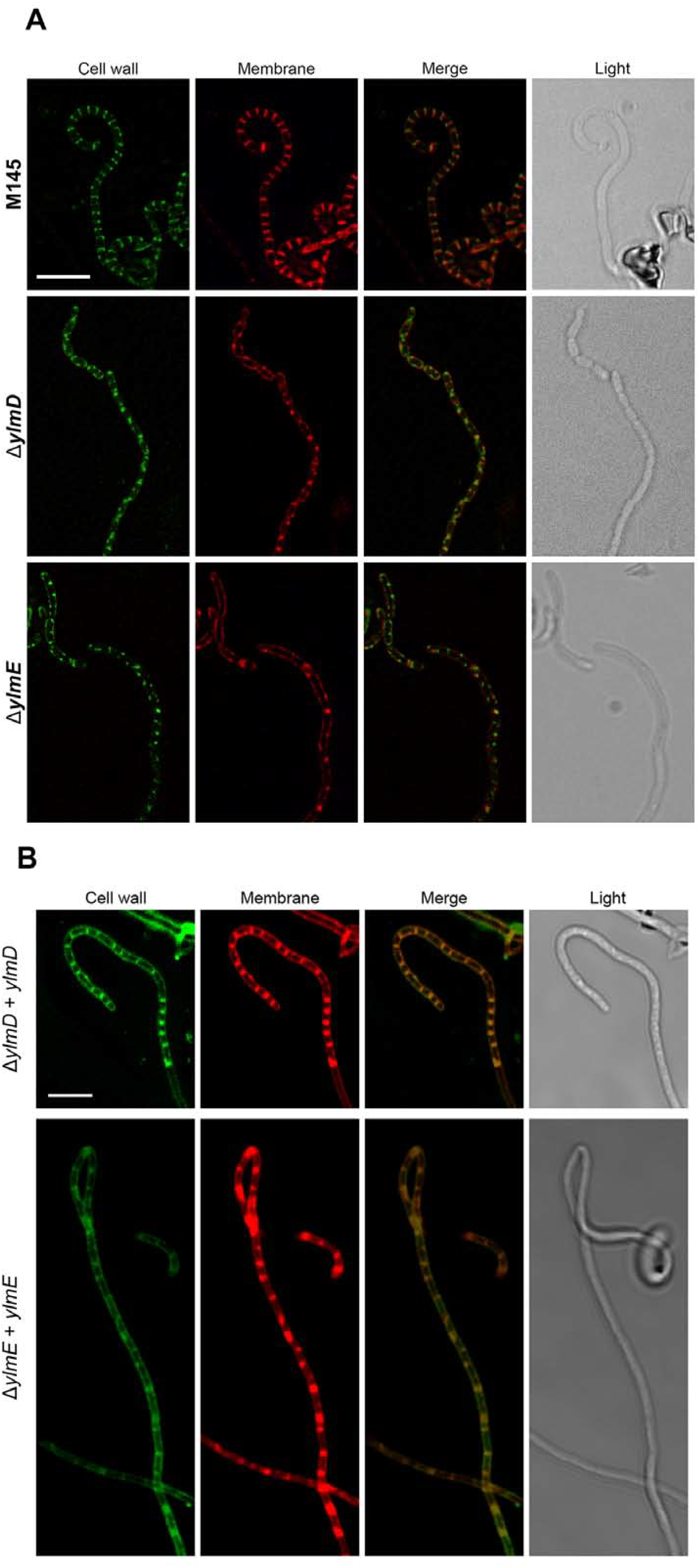
Fluorescence microscopy of cell wall, DNA and membranes. (A) Fluorescent micrographs of hyphae stained for cell-wall synthesis (FITC-WGA or Oregon-WGA) or membranes (FM5-95). An overlay of these images is presented in the third column, and the corresponding light image in the last column. Bar, 5 μm. (B) Fluorescence micrographs showing DNA and cell-wall distribution in the complemented *ylmD* and *ylmE* mutants. While ladders of septa were formed in both strains, indicative that sporulation was restored to the mutants, in particular the complemented *ylmE* mutant formed imperfect septa. Cultures were grown on SFM agar plates for 5 days at 30°C. Bar, 10 μm.

### Cellular localization of YlmD and YlmE

Given the impact of *ylmDE* on sporulation, we analyzed how YlmD and YlmE were localized in the hyphae of *S. coelicolor*. To this end, constructs based on the low-copy number vector pHJL401 were prepared to allow the expression of YlmD-eGFP and YlmE-eGFP fusion proteins, which were expressed from the natural *ftsZ* promoter region (see Materials and Methods section for details). These constructs were then introduced into *S. coelicolor* M145 and (as a control for functionality) in the mutants. The constructs partially restored development to the respective mutants, with significant restoration of the sporulation defects in *ylmD* mutants, and partial restoration of sporulation to *ylmE* mutants (Fig. 2B).

In vegetative hyphae, only very weak signals were obtained for YlmD-eGFP and YlmE-eGFP, indicative of low protein expression (Fig. 6A). In early and late aerial hyphae, both YlmD-eGFP and YlmE-eGFP became more visible and showed diffuse localization along the wall of the aerial hyphae (Fig. 6A). During early sporulation, YlmD-eGFP and YlmE-eGFP showed an irregular pattern of varying intensity, with some spores showing bright fluorescence indicative of high concentrations of YlmD or YlmE, while others hardly showed any fluorescence (Fig 6A). This suggests that YlmD and YlmE are differentially expressed throughout the nascent spore chains, and do not exclusively co-localize with FtsZ and the divisome. To rule out possible proteolysis of YlmD- or YlmE-eGFP fusion proteins, Western analysis was performed on extracts of surface-grown *S. coelicolor* mycelia, using antibodies against GFP. *S. coelicolor* M145 with empty vector and *S coelicolor* M145 expressing freely mobile eGFP ^41^ were included as controls. Only a single band was identified in strains GAL50 and GAL49, corresponding to the expected lengths for YlmD-eGFP and YlmE-eGFP, respectively (Fig. S3). In both cases, the predicted mass of the fusion protein is around 52 kDa, which conforms well to the apparent mobility seen by Western analysis. The strain expressing only eGFP showed the expected band of around 27 kDa for the non-fused protein. Therefore, we conclude that the observed localizations can indeed be ascribed to intact YlmD- or YlmE-eGFP.

**Figure 6.**
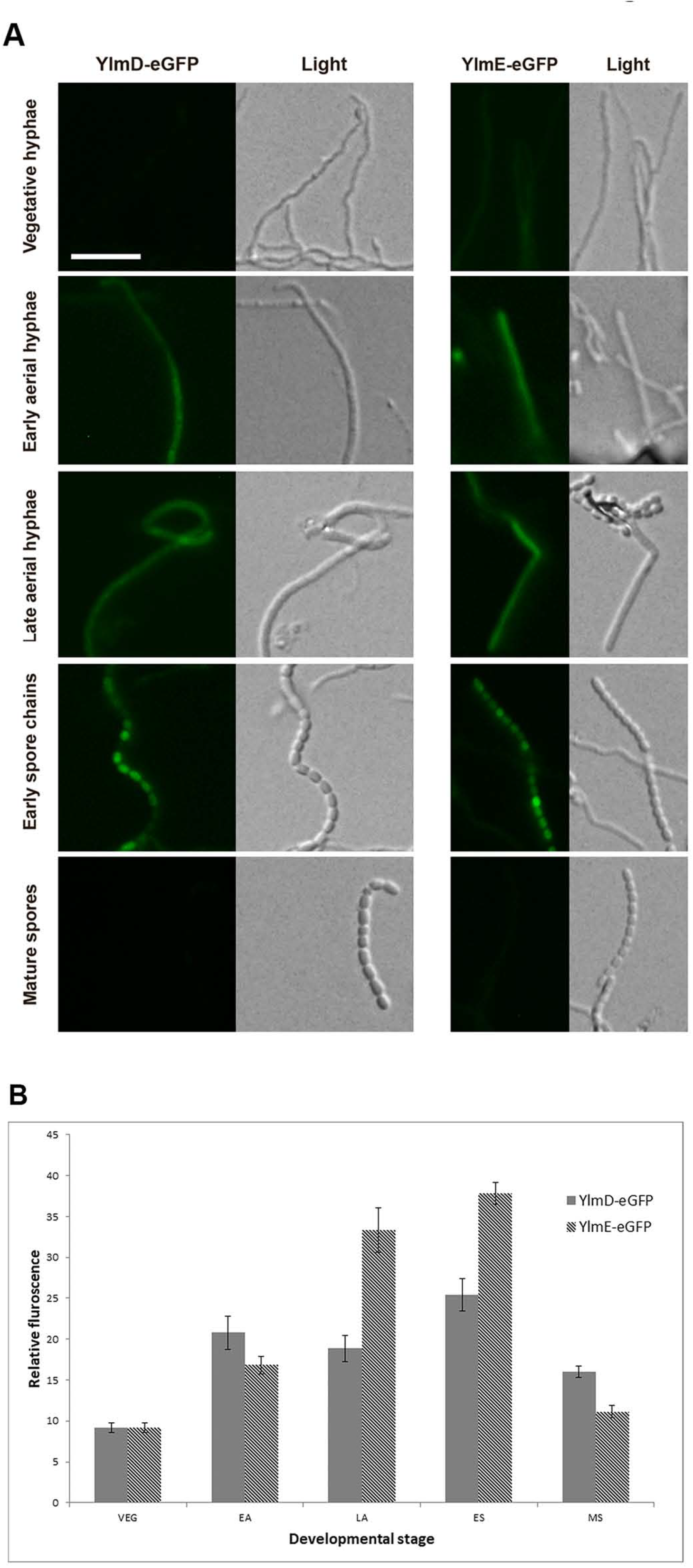
Localization of YlmD-GFP and YlmE-GFP. (A) Sporogenic aerial hyphae of *S. coelicolor* M145 at different stages of development were imaged by fluorescence microscopy visualizing YlmD-eGFP and YlmE-eGFP. Stages were: vegetative growth, early aerial development, late aerial development, early sporulation and mature spores. During spore maturation, YlmD and YlmE had a ‘patchy’ localization, suggesting that at this stage YlmD and YlmE may co-localize with the cell-wall synthetic machinery. Bar, 5 μm. (B) Relative fluorescence intensity of YlmD-eGFP and YlmE-eGFP during *Streptomyces* development. VEG, vegetative growth; EA, early aerial growth; LA, late aerial growth; ES, early sporulation; MS, mature spores.

The fluorescence intensity of YlmD-eGFP and YlmE-eGFP was measured on images representing different developmental stages (Fig. 6B). As development progressed, the fluorescence intensities increased for both YlmD-eGFP and YlmE-eGFP and reached peak levels during the phase corresponding to late aerial growth and early sporulation. When spores matured, YlmD-eGFP and YlmE-eGFP signals decreased to the level as in vegetative growth. These results suggest YlmD and YlmE proteins are active primarily during the phase of sporulation-specific cell division.

### Localization of FtsZ is disturbed in *ylmD* and *ylmE* null mutants

To investigate the localization of FtsZ in the mutants, integrative plasmid pKF41 that expresses FtsZ-eGFP from the native *ftsZ* promoter region ^42^ was introduced into the *ylmD* and *ylmE* mutants, to generate strain GAL52 and GAL53, respectively. Prior to sporulation, typical ladders of Z-rings were observed in the parental strain *S. coelicolor* M145, while *ylmD* and *ylmE* null mutants showed abnormal Z-ladders (Fig. 7). In the absence of YlmD, ladders were still observed, but the intensity varied and spacing between the individual rings was less regular, with many neighboring Z-rings either close together or widely spaced. Consistent with the sporulation defect, *ylmE* null mutants produced very few Z-rings, and the few Z-ladders that where formed were highly irregular or unfinished, and significantly shorter than in the parental strain. Thus, FtsZ localization is irregular in *ylmD* mutants and highly compromised in *ylmE* mutants.

**Figure 7.**
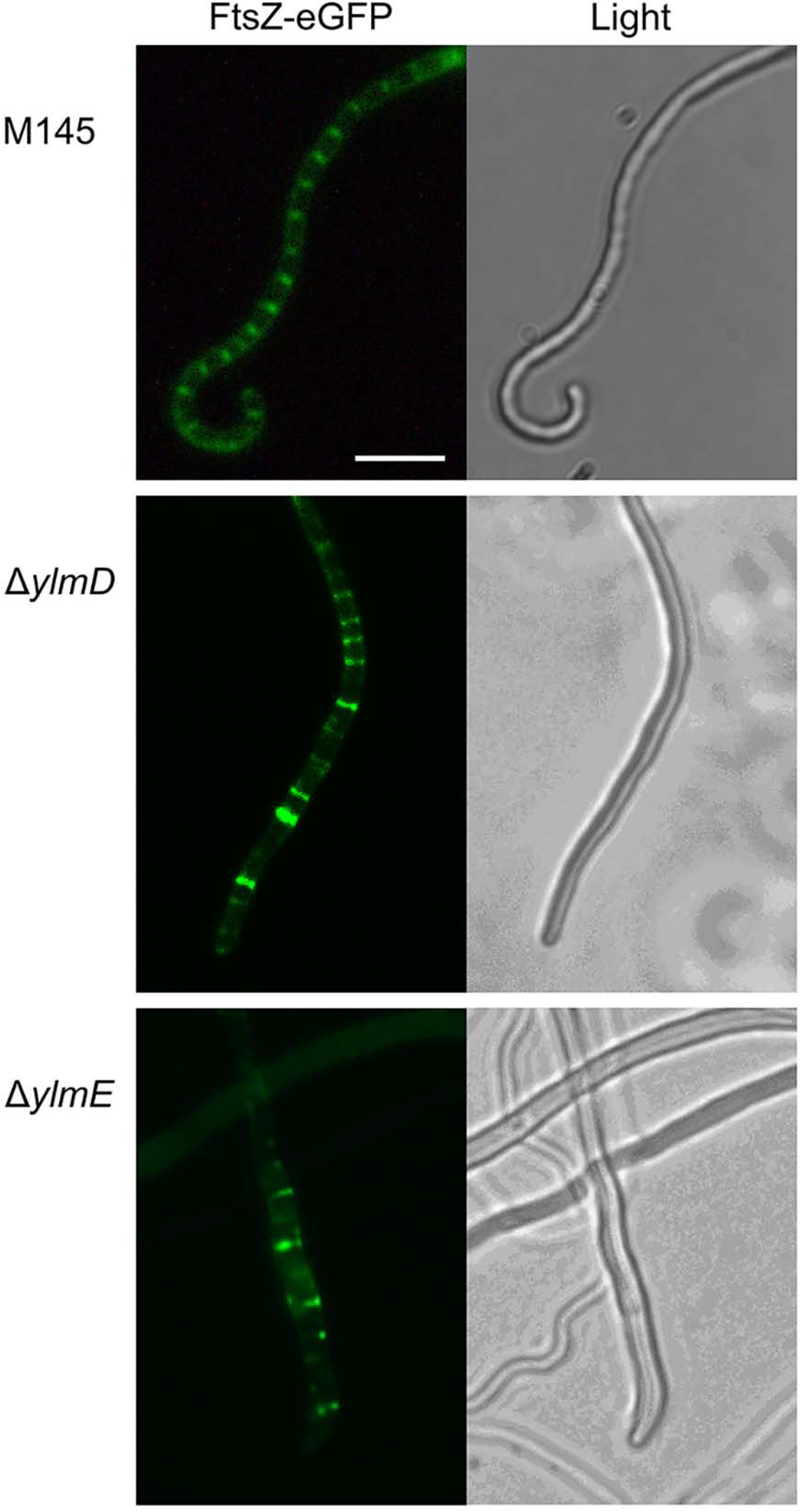
Localization of FtsZ-eGFP in *S. coelicolor* M145 and its *ylmD* and *ylmE* mutants. FtsZ-eGFP formed typical ladders in wild-type cells (M145). In contrast, YlmE is required for the formation of ladders of FtsZ, while the absence of YlmD caused irregular spacing between the septa. Cultures were grown on SFM agar plates for 5 days at 30°C. Bar, 5 μm.

### A model for sporulation control by YlmDE in streptomycetes

Our data show that mutants of *S. coelicolor* lacking *ylmE* have severe developmental defects, with highly compromised cell-wall synthesis and aberrant cell division. Mutants which lack *ylmD* sporulated well, though many spores exhibited aberrant shapes and variable lengths. Surprisingly, the additional deletion of *ylmD* greatly rescued the sporulation-deficient phenotype of *ylmE* mutants, with the *ylmDE* null mutant producing abundant and mostly regularly sized spores.

We propose that *YLMD* AND *YLME* Form a two-part system, reminiscent of toxin-antitoxin systems ^43^. Similarly dysfunction of one protein in the presence of another has also been reported for the sporulation control protein WhiJ, following the observation that media-dependent sporulation defects caused by a point mutation in *whiJ* (SCO4543) could be rescued by the full deletion of *whiJ* ^44^. The authors proposed a model wherein repression of developmental genes by WhiJ could be released when WhiJ interacts with a WhiJ-associated protein encoded by the adjacent gene SCO4542 ^44^. It was suggested that the aberrant protein WhiJ* still binds to operator sequences of developmental genes, leading to their transcriptional repression, but fails to associate with SCO4542, with a permanent block of transcription of developmental genes as a result ^44^. Similarly, the expression of YlmD alone is detrimental for sporulation, suggesting that YlmD acts in a deleterious manner towards sporulation, yet this effect can be relieved by the presence of YlmE. Supporting evidence for this is that deletion of both genes allows *Streptomyces* to develop normally, and by the observation that *ylmD* is never found alone in bacterial genomes, while the occurrence of an orphan *ylmE* is seen frequently. Further support comes from the report that *ylmDE* double mutants of *S. venezuelae* also sporulate relatively normally ^32^. This also indicates that this two-part system functions in a similar manner in a range of *Streptomyces* species. This apparently rules out major divergence between the two morphologically distinct *Streptomyces* clades, namely those that sporulate in submerged cultures and those that do not ^45,46^. However, it will be very interesting to see if submerged sporulation of *S. venezuelae* is prevented by the deletion of just *ylmE*. Currently, we are performing detailed structural and functional analysis of YlmD and YlmE, to elucidate their biochemical function and their precise role in bacterial cell division.

## METHODS

### Phylogenetic analysis of *ylmD* and *ylmE*

The amino acid sequence of YlmD and YlmE were extracted from StrepDB (http://strepdb.streptomyces.org.uk) and used to search the NCBI database (www.ncbi.nlm.nih.gov) using BLASTP against the non-redundant protein sequence database. Alignment of YlmD and YlmE was generated using ClustalW ^47^ followed by manual editing in MEGA v. 4.0. The neighbour-joining trees ^48^ were generated with default parameters settings as implemented in MEGA v. 4.0 ^49^. The maximum-likelihood trees were made using the best fit models predicted by MEGA. Tree reliability was estimated by bootstrapping with 1000 replicates. Trees were drawn with either N-J or ML algorithms gave trees of similar topologies indicating that the phylogenies are likely to reliable.

### Bacterial strains and media

All bacterial strains used in this study are listed in Table S1. *E. coli* JM109 was used for routine cloning and ET12567 ^50^ to prepare nonmethylated DNA to bypass the methyl-specific restriction system of *S. coelicolor. E. coli* strains were propagated in Luria broth, where appropriate supplemented with antibiotics for selection, namely ampicillin (100 μg/ml end concentration), apramycin (50 μg/ml) and/or chloramphenicol (25 μg/ml). *S. coelicolor* A3(2) M145 ^51^ and *S. lividans* 66 ^52^ were obtained from the John Innes Centre strain collection. *S. coelicolor* strains were grown on soya flour medium (SFM) or minimal media mannitol (MM) agar plates for phenotypic characterization and on R5 agar plates for regeneration of protoplasts ^53^. Antibiotics used for screening Streptomyces were apramycin (20 μg/ml end concentration) and thiostrepton (10 μg/ml).

### Plasmids and constructs

All plasmids and constructs described in this study are summarized in Table S2. The oligonucleotides used for PCR are listed in Table S3. PCR reactions were performed using Pfu DNA polymerase as described ^54^. All inserts of the constructs were verified by DNA sequencing, which was performed at BaseClear (Leiden, The Netherlands).

#### Constructs for the deletion of ylmD and ylmE

The strategy for creating knock-out mutants is based on the unstable multi-copy vector pWHM3 ^55^ as described previously ^56^. For each knock-out construct roughly 1.5 kb of upstream and downstream region of the respective genes were amplified by PCR from cosmid St4A10 that contains the *dcw* cluster of *S. coelicolor*. The upstream region was thereby cloned as an EcoRI-XbaI fragment, and the downstream part as an XbaI-BamHI fragment, and these were ligated into EcoRI-BamHI-digested pWHM3 (for the precise location of the oligonucleotides see Table S3). In this way, an XbaI site was engineered in-between the flanking regions of the gene of interest. This was then used to insert the apramycin resistance cassette *aac(3)IV* flanked by *loxP* sites, using engineered XbaI sites. The presence of the *loxP* recognition sites allows the efficient removal of the apramycin resistance cassette following the introduction of a plasmid pUWL-Cre expressing the Cre recombinase ^57,58^. Knock-out plasmids pGWS728 and pGWS729 were created for the deletion of nucleotide positions +25/+696 of *ylmE* (SCO2080) and +25/+705 of *ylmD* (SCO2081), whereby +1 refers to the translational start site of the respective genes. This allowed first the replacement by the apramycin resistance cassette. Subsequently the apramycin resistance cassette was removed using expression of pUWLCre. To create a *ylmDE* double deletion mutant, construct pGWS1044 was created which contains the upstream region of *ylmD* and downstream region of *ylmE* flanked by *loxP* sites, and with the apramycin resistance cassette *aac(3)IV* inserted in between.

For complementation of the *ylmE* and *ylmD* null mutants, the entire coding regions of SCO2080 and SCO2081 (with stop codons) were amplified from the *S. coelicolor* M145 chromosome using primer pairs ylmE_F+1 and ylmE_R+723 and ylmD_F+1 and ylmD_R+732, respectively. The PCR products were digested with StuI/XbaI, and inserted downstream of the native *ftsZ* promoter region in pHJL401, respectively. Thus constructs pGWS1042 and pGWS1043 were generated that express *ylmE* and *ylmD*, respectively, under control of the *S. coelicolor ftsZ* promoter region. Alternatively, construct pKR8 was used for complementation; pKR8 is based on integrative vector pIJ8600 ^59^ and contains the 2227742-2241015 region of the *S. coelicolor* genome, encompassing the end of *murX* (SCO2087), the entire coding sequences of *murD, ftsW, murG, ftsQ, ftsZ, ylmD, ylmE, sepF, sepG, divIVA* and a large part of SCO2076 (encoding Ile-tRNA synthetase).

#### Constructs for the localization of YlmD and YlmE

The entire coding regions of SCO2080 and SCO2081 (without stop codons) were amplified from the *S. coelicolor* M145 chromosome using primer pairs ylmE_F+1 and ylmE_R+717 and ylmD_F+1 and ylmD_R+726, respectively. The PCR products were digested with StuI/BamHI, and inserted downstream of the native *ftsZ* promoter region and immediately upstream of *egfp* in pHJL401. The latter is a highly stable vector with low copy number that generally results in wild-type transcription levels and is well suited for among others complementation experiments ^40^. Thus constructs pGWS757 and pGWS758 were generated that express YlmE-eGFP and YlmD-eGFP, respectively, from the native *S. coelicolor ftsZ* promoter region. Plasmid pKF41 expresses FtsZ-eGFP from its own promoter region ^42^. Constructs pGWS757 and pGWS758 were also used to complement *ylmE* and *ylmD* null mutants, respectively.

### eGFP detection using western blot analysis

Strains were grown on SFM agar plates containing the appropriate antibiotics overlaid with cellophane disks. Biomass (mycelium and spores) was collected after 6 days of growth at 30 °C and suspended in urea lysis buffer (8M urea, 2M thiourea, 4% CHAPS, 1mM PMSF and 65mM DTT), followed by cell disruption via sonication. Western blots were performed as described ^60^. Briefly, cell lysates were obtained after centrifugation at 30,000x g for 30 min at 4°C. Proteins were separated on a 12.5% SDS-PAGE gel, and then transferred to PVDF membrane using Mini Trans-Blot (Bio-Rad). As primary antibody a 1:2000 dilution of GFP antibody (ThermoFisher, REF A11122) was used and incubated for 16 h at 4°C in the presence of 3% BSA. After washing, the membrane was incubated with alkaline phosphatase conjugated anti-rabbit IgG (Sigma) as secondary antibody for 1 h at room temperature. Visualization was done using BCIP/NBT color development substrate.

### Microscopy

Fluorescence and light microscopy were performed as described previously ^61^. For peptidoglycan staining we used FITC-labeled wheat germ agglutinin (FITC-WGA) or Oregon Green 488 conjugated WGA (Oregon-WGA); for membrane staining, we used FM5-95 (all obtained from Molecular Probes). All images were background-corrected, setting the signal outside the hyphae to 0 to obtain a sufficiently dark background. These corrections were made using Adobe Photoshop CS4. Cryo-scanning electron microscopy (cryo-SEM) and transmission electron microscopy (TEM) were performed as described ^62^. For quantification of fluorescence intensity of YlmD-eGFP and YlmE-eGFP, the arbitrary average intensity values of manually selected areas of interest were determined as relative fluorescence intensity. All green fluorescent images were made with equal exposure time.

### Data analysis

For the Kolmogorov–Smirnov (KS) test, the critical value *D_crit_* was calculated for the number of measured spores (n) and significance level (α) of 0.05: 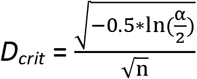. The difference (*D_n_*) between the observed cumulative distribution function (*F_obs_*) and the expected cumulative distribution function (*F_exp_*) was calculated as: *D_n_* = |*F_exp_(x*) − *F_obs_(x*)|. The maximal value was identified as *D_max_*. The null hypothesis that the data comes from the specified distribution was rejected if *D_max_* > *D*_crit_. The Mann-Whitney U test was done using the SPSS software release 24.0.

### Computer analysis

DNA and protein database searches were completed by using StrepDB (http://strepdb.streptomyces.org.uk/). Phylogenetic analysis was done using the STRING engine at EMBL (www.string.EMBL).

## DATA AVAILABILITY

All strains, materials and data will be made available upon request.

## ACKNOWLEDGEMENTS

This work was supported by the Netherlands Organization for Scientific Research (NWO), via VICI grant 10379 to GPvW.

## COMPETING FINANCIAL INTERESTS

The authors declare no conflict of interests.

## AUTHOR CONTRIBUTIONS

LZ performed the molecular biology experiments and created the mutants and expression constructs, JW performed the imaging and PAH the phylogenetic analysis. GVW conceived the experiments together with the other authors. All authors wrote and approved the manuscript.

